# Imaging through Wind*an*see electrode arrays reveals a small fraction of local neurons following surface MUA

**DOI:** 10.1101/2022.09.01.506113

**Authors:** Martin Thunemann, Lorraine Hossain, Torbjørn V. Ness, Nicholas Rogers, Keundong Lee, Sang Heon Lee, Kıvılcım Kılıç, Hongseok Oh, Michael N. Economo, Vikash Gilja, Gaute T. Einevoll, Shadi A. Dayeh, Anna Devor

## Abstract

Prior studies have shown that neuronal spikes can be recorded with microelectrode arrays placed on the cortical surface. However, the etiology of these spikes remains unclear. Because the top cortical layer (layer 1) contains very few neuronal cell bodies, it has been proposed that these spikes originate from neurons with cell bodies in layer 2. To address this question, we combined two-photon calcium imaging with electrophysiological recordings from the cortical surface in awake mice using chronically implanted PEDOT:PSS electrode arrays on transparent parylene C substrate.

Our electrode arrays (termed Wind*an*see) were integrated with cortical *wind*ows offering *see*-through optical access while also providing measurements of local field potentials (LFP) and multiunit activity (MUA) from the cortical surface. To enable longitudinal data acquisition, we have developed a mechanical solution for installation, connectorization, and protection of Wind*an*see devices aiming for an unhindered access for high numerical aperture microscope objectives and a lifetime of several months while worn by a mouse.

Contrary to the common notion, our measurements revealed that only a small fraction of layer 2 neurons from the sampled pool (~13%) faithfully followed MUA recorded from the surface above the imaging field-of-view. Surprised by this result, we turned to computational modeling for an alternative explanation of the MUA signal. Using realistic modeling of neurons with back-propagating dendritic properties, we computed the extracellular action potential at the cortical surface due to firing of local cortical neurons and compared the result to that due to axonal inputs to layer 1. Assuming the literature values for the cell/axon density and firing rates, our modeling results show that surface MUA due to axonal inputs is over an order of magnitude larger than that due to firing of layer 2 pyramidal neurons.

Thus, a combination of surface MUA recordings with two-photon calcium imaging can provide complementary information about the input to a cortical column and the local circuit response. Cortical layer I plays an important role in integration of a broad range of cortico-cortical, thalamocortical and neuromodulatory inputs. Therefore, detecting their activity as MUA while combining electrode recording with two-photon imaging using optically transparent surface electrode arrays would facilitate studies of the input/output relationship in cortical circuits, inform computational circuit models, and improve the accuracy of the next generation brain-machine interfaces.

## Introduction

With the advent of genetically encoded calcium probes of neuronal activity (Dana et al 2019, Qian et al 2020), computational methods for extraction of spikes from calcium signals (Giovannucci et al 2019), microscopic tools for large-scale imaging (reviewed in (Abdelfattah et al 2022)) and electrode array technology (Hong & Lieber 2019, Lee et al, Vazquez-Guardado et al 2020), there has been a growing appreciation of the need to reconcile optical and electrophysiological measurements of neuronal spikes (Siegle et al 2021). Among the novel electrode technologies are minimally invasive devices that can be placed on the cortical surface to record extracellular electrical potentials, which can be then separated into high-frequency spiking activity and low-frequency “brain waves” (Buzsaki et al 2015, Choi et al 2019, Ganji et al 2019, Hermiz et al 2020, Hong & Lieber 2019, Jun et al 2017, Khodagholy et al 2015, Paulk et al 2021, Tchoe et al 2022). Because the top layer of cerebral cortex (layer 1, L1) has low density of neuronal cell bodies, spiking activity picked up from the cortical surface has been attributed to neurons with cell bodies located in layer 2/3 (L2/3; ~100 μm deep in the mouse) (Khodagholy et al 2015). In the present work, we sought to evaluate this hypothesis using two-photon calcium imaging through an optically transparent surface electrode array.

Most space-resolved neurorecording devices are incompatible with optical imaging because electrode arrays are not transparent, and electronic circuits are located immediately above the electrodes covering the brain surface blocking the visibility and/or access for optical instruments such as microscope objectives. Two general solutions have been proposed to allow simultaneous optical imaging. The first one leverages high transparency of monolayer graphene. Microelectrode arrays made of graphene or carbon nanotubes on transparent parylene C or other substrates have been employed in combination with single- and multiphoton optical imaging *in vivo* by us and others (Driscoll et al 2021, Kuzum et al 2014, Lu et al 2018, Park et al 2016, Park et al 2014, Thunemann et al 2018, Zhang et al 2018). The second strategy is using non-transparent metal and polymer electrodes on a transparent substrate, where the electrodes are small enough not to produce significant shadows (Donahue et al 2018, Donaldson et al 2022, Ganji et al 2018, Ganji et al 2019, Hossain et al 2020, Qiang et al 2018, Seo et al 2019). Following the second strategy, we have previously developed stable and biocompatible Poly(3,4-ethylenedioxythiophene)-poly(styrenesulfonate) (PEDOT:PSS) surface electrode arrays suitable for chronic implantation (Ganji et al 2018, Ganji et al 2019, Hossain et al 2020). In the present study, we leveraged this advance to engineer wearable devices for seamless integration of electrical and optical measurements of cortical neuronal activity in awake mice. We coined our wearable devices, which replicate the functionality of conventional cranial *wind*ows *an*d offer *see*-through optical access, Wind*an*see after the famous Windansea Beach in La Jolla, California.

Using chronically implanted Wind*an*see devices and combining electrophysiological recording with two-photon calcium imaging, we show that spiking of local cortical neurons is unlikely to fully account for multiunit activity (MUA) recorded at the cortical surface. This is because only a small fraction of these neurons exhibited calcium transients synchronous with the MUA signal. Another potential contributor to surface MUA is axons projecting to L1. To evaluate this possibility, we simulated the extracellular potential at the cortical surface due to (i) spiking of realistic cortical excitatory neurons across layers, and (ii) axonal afferents in L1. Assuming the literature values for the cell/axon density and firing rates, our modeling results revealed that axonal inputs generated surface MUA signals were an order of magnitude larger compared to those generated by firing of local L2/3 neurons.

Cortical L1 plays an important role in integration of a broad range of cortico-cortical, thalamocortical and neuromodulatory inputs. Therefore, detecting their activity as MUA while combining electrode recording with two-photon imaging using optically transparent surface electrode arrays would facilitate studies of the input/output relationship in cortical circuits, inform computational circuit models, and improve the accuracy of the next generation brain-machine interfaces.

## Methods

### Fabrication and characterization of PEDOT:PSS microelectrode arrays

We fabricated 32-channel PEDOT:PSS/parylene C microelectrode arrays with electrodes arranged in square grids with 0.2 mm spacing and 20 μm contact (electrode) diameter according to previously published protocols (Ganji et al 2018, Ganji et al 2019, Hossain et al 2020). Briefly, after spin-coating of an anti-adhesion layer, parylene C was deposited on the carrier wafer to a thickness of 2.1-2.5 μm. Metallization of the electrodes and leads was conducted via electron beam evaporation with a thickness of 10-nm Ti and 100-nm Au. A second 2.1-2.5-μm thick layer of parylene C was deposited, followed by spin-coating of an anti-adhesion layer and deposition of a third, sacrificial parylene C layer. A subsequent layer of SU-8 was used as a hard mask for parylene C dry etching, leaving contacts and connector pads exposed and the remaining metal leads encapsulated. After coating with PEDOT:PSS solution, the sacrificial parylene C layer was peeled off, and thermal curing was performed. The final device was manually cut and separated from the hard carrier wafer resulting in the microelectrode array. Microelectrode arrays for acute experiments were bonded to ribbon cables using conductive tape (Elform Heat Seal Connectors, Yutek Tronic). For chronically implanted microelectrode arrays, the connector was bonded to a miniature printed circuit board (PCB) with conductive epoxy (Silver Conductive Epoxy Adhesive, MG Chemicals) and cured at room temperature within a vacuum desiccator.

Electrophysiological properties of the microelectrode arrays were characterized with electrochemical impedance spectroscopy (EIS) using a conventional three-terminal setup in phosphate-buffered saline (PBS) solution, with Ag/AgCl as the reference electrode, Pt as the counter electrode, and the PEDOT:PSS device as the working electrode.

### Animals

All experiments were conducted in accordance with the National Institutes of Health’s Guide for the Care and Use of Laboratory Animals and were approved by the Institutional Animal Care and Use Committee (IACUC) at the University of California San Diego and Boston University. We used 12 adult (age >12 weeks) mice of either sex including three ICR mice (bred in house), two GAD67-EGFP mice on ICR background (Tamamaki et al 2003), four C57Bl/6J (The Jackson Laboratories, Stock 000664), and three Emx1-Cre/Ai32 mice; the latter were bred in-house from the parental strains (The Jackson Laboratories, stocks 005628 and 024109). Animals were kept under standard conditions in individually ventilated cages on a 12-h light/dark cycle with *ad libitum* access to food and water.

### Surgical procedures for acute experiments

Surgical procedures were performed as described previously (Rogers et al 2019, Thunemann et al 2018) using ICR or GAD67-EGFP mice on ICR background weighing 25-35 g. Briefly, the mouse was placed on a heating pad and anesthetized with isoflurane in oxygen (5% for induction, 1-2% for maintenance). The left femoral artery was catheterized for blood pressure monitoring and drug/dye injection, and a tracheotomy was performed for mechanical ventilation. After fixing the skull to a metal holder with dental acrylic, craniotomy and durotomy were performed over the right whisker-barrel and surrounding cortex. A well was formed around the craniotomy using dental acrylic, and the exposure was filled with artificial CSF (ACSF). During placement of the microelectrode array, the brain surface was dried briefly to promote contact of the electrodes with the cortical surface. Following placement, the array and surrounding cortex were covered with 0.7% agarose in ACSF.

In experiments involving validation of optical transparency, Sulforhodamine 101 (SR101) dissolved in ACSF was applied to the surface prior to placement of the array for ca. 60 s; excess dye was removed with several washes with fresh ACSF. A drop of agarose in ACSF was applied on the brain surface, and the exposure was covered with a glass coverslip and sealed with dental acrylic.

### Surgical procedures for chronic experiments

Surgical procedures were performed as previously described (Desjardins et al 2019, Kilic et al 2020). Briefly, dexamethasone (4.8 mg/kg) was injected ~4 h prior to surgery. Mice were anesthetized with isoflurane and secured in a stereotaxic frame. The skull was exposed bilaterally by removing the skin over an area of ~1.5 cm x 1.5 cm. Then, the headpost was attached to the bone with dental resin. A steel screw (#000) was inserted into the bone above the cerebellum. The bone along a ~3.5-mm circumference (center coordinates A-P 2 mm and L-R 3 mm relative to Bregma) and the underlying dura were removed. 250-500 nL containing 3×10^12^ GC/mL of the recombinant adeno-associated virus pAAV.Syn.NES-jRGECO1a.WPRE.SV40 (Dana et al 2016), a gift from Douglas Kim & GENIE Project (Addgene viral prep #100854-AAV9), was injected at 2-3 sites 300-400 μm below the cortical surface using a microinjector (Nanoject III, Drummond). After placing the Wind*an*see device within the exposure, the circumference of the 5-mm glass was sealed and fixed to the bone with dental resin. Buprenorphine (0.05 mg/kg) was injected before discontinuation of anesthesia. A combination of 0.53 mg/mL sulfamethoxazole and 0.11 mg/mL trimethoprim (Sulfatrim), and ibuprofen (0.05 mg/mL) was provided with the drinking water starting on the day of surgery and for five days after surgery. Full recovery and return to normal behavior were generally observed within 48 h after surgery.

### Structural two-photon imaging

We used two-photon microscopy to validate suitability of PEDOT:PSS electrode arrays for deep, high-resolution optical imaging. To visualize the vasculature, we used Alexa Fluor 680 conjugated to 2-MDa Dextran (Kobat et al 2009, Li et al 2019) produced in-house. Images were obtained using an Ultima two-photon laser scanning microscopy system from Bruker Fluorescence Microscopy. EGFP and SR101 were excited at 920 nm using an Ultra II femtosecond Ti:Sapphire laser (Coherent), while Alexa 680-Dextran was excited at 1240 nm using an optical parametric oscillator (Chameleon Compact OPO, Coherent) pumped by the same Ti:Sapphire laser. We used cooled GaAsP detectors (Hamamatsu, H7422P-40) or multi-alkaline photomultiplier tubes (Hamamatsu, R3896) for signal detection in combination with the following emission filters: EGFP, 525/50 nm, SR101, 617/73 nm, Alexa 680, 795/150 nm. We used a 4x objective (Olympus XLFluor4x/340, NA=0.28) to obtain low-resolution images of the cranial exposure. A 0.5-NA 20x water-immersion objective (Olympus UMPlanFI) was used for high-resolution imaging.

### Electrophysiological recordings under anesthesia

Prior to recording, the mouse was paralyzed with pancuronium bromide (0.4 mg/kg/h IV, P1918, Sigma) and artificially ventilated (~110 min^-1^); anesthesia was switched to α-chloralose (50 mg/kg/h IV, C0128, Sigma or 100459, MP Biochemicals) prior to data acquisition. A tungsten extracellular microelectrode (FHC, 6-8 MΩ) was used to determine the location of the C1 whisker representation on the whisker-barrel cortex prior to electrode array placement. The reference electrode was an Ag/AgCl ball placed next to the skull. Single whiskers were deflected upward by a wire loop coupled to a computer-controlled piezoelectric stimulator using 2-s interstimulus interval; recordings included spontaneous epochs as well as periods with stimulation. The PEDOT:PSS electrode array was connected to a RHD2000 amplifier board and RHD2000 evaluation system (Intan Technologies) using a custom-build connector. Broadband electrophysiological data above 0.1 Hz were acquired at 20 kHz.

### Simultaneous electrophysiological and optical measurements in chronically implanted mice

Starting at least seven days after the surgical procedure, mice were habituated in one session per day to accept increasingly longer periods (up to 2 h) of head restraint under the microscope objective. During head restraint, the mouse was placed on a suspended bed and rewarded with sweetened condensed milk every 15-20 min. At the beginning of each recording session, mice were briefly anesthetized with isoflurane in oxygen (5% induction, 1-1.5% maintenance) to perform head fixation and establish connection to the recording setup. A ribbon cable connected the implanted PCB to a secondary PCB equipped with a dual row horizontal Nano Strip connector (Omnetics Connector Corporation) and ground and reference inputs. Miniature alligator clips were used to connect the reference screw to the PCB board, and the ground input of the device was connected to the metal stage, which was connected through the microscope table to a common ground. Then, anesthesia was removed, and the mouse was allowed to recover for 15-20 min before starting data acquisition. Electrophysiological data were acquired as in acute experiments.

Two-photon calcium imaging was performed using the same Ultima two-photon laser scanning microscopy system used for structural imaging. Calcium biosensor jRGECOa1 was excited at 1100 nm using the OPO. Emitted light was directed to GaAsP detector (Hamamatsu, H7422P-40) though a bandpass 617/73-nm filter. Rectangular fields-of-view (FOVs) with a size of ~130×110 μm were imaged in frame-scan mode at the target acquisition rate of ~15 Hz.

Sensory stimulus consisted of a brief air puff delivered to the lower bottom part of the contralateral whisker pad (to avoid an eye blink) using a plastic tube connected to a pneumatic pump (PV820 Pneumatic PicoPump, WPI). We used 30 trials per run with 5-s inter-stimulus interval, ISI.

### Motion detection

We used a CCD camera (acA1920-150um, Basler) attached to a variable zoom lens (Navitar 7000 Macro Lens) and a 940-nm LED (M940L3, Thorlabs) to monitor movement of awake head-fixed mice. During electrophysiological recording without calcium imaging, camera and LED were operated in a free-running mode; an additional broad-spectrum LED within the camera’s field of view was used for offline synchronization of the electrophysiological data with the mouse video and stimulus onset. During two-photon imaging, the CCD camera and 940-nm LED were triggered by the end-of-frame trigger, such that an image of the mouse was acquired in-between the acquisition of individual frames in two-photon time series. To extract motion from the recorded movies, temporal variance of pixel brightness within a user-defined region of interest, which included the mouse face and forelimbs, was estimated using a moving window. A manually defined threshold was imposed to identify periods of the animal movement.

### Synchronization of two-photon imaging, electrophysiological recordings, and sensory stimulation

Synchronization of two-photon acquisition, electrophysiological recordings, and stimulus delivery was achieved using a dedicated computer equipped with a multifunction input/output (I/O) data acquisition (DAQ) card (NI-6229, National Instruments) controlled with a custom-written MATLAB script. The same DAQ was used to (1) trigger the sensory stimulus and (2) record the CCD and two-photon frame transistor–transistor logic (TTL) signals for offline synchronization during data analysis. In addition, TTL signals generated by the DAQ card were recorded by the Intan setup.

### Data analysis

Electrophysiological data were analyzed in MATLAB. Independent component analysis (ICA) was performed using the *jadeR* function adapted from the publicly available MATLAB-based EEGLab resource (https://eeglab.org/) (Delorme & Makeig 2004) to mitigate movement-related artifacts in the electrophysiological data. We removed up to three independent components that had unrealistically high amplitude while lacking spatial heterogeneity or followed the time-course of laser scanning.

Raw electrophysiological data were low-pass filtered at 250 Hz and resampled to 4000 Hz to isolate the local field potential (LFP). Multiunit activity (MUA) was obtained by high-pass filtering above 350 Hz. To calculate the MUA envelope (eMUA), MUA signals were temporally smoothed using a Gaussian kernel (FWHM=50 ms).

Frequency power analysis of the low-pass filtered data was performed with Morlet wavelets (log_10_(0.05 Hz) – log_10_(180 Hz) in 100 logarithmic steps) using the ‘morlet_transform’ function (fc = 1 Hz, FWHM = 2 s) of the MATLAB-based BrainStorm3 environment (Tadel et al 2011). To estimate the frequency power of common frequency bands (δ, <4 Hz; θ, 4-8 Hz; α, 8-12 Hz, β, 12-30 Hz; γ, 30-80 Hz), wavelet coefficients of the respective frequency range were averaged. For display, individual frequency bands were isolated from the LFP signal for display using lowpass (δ band) or bandpass (θ, α, β, γ bands) filters.

Data from two-photon calcium imaging were analyzed in MATLAB using custom-written software. Regions of interest (ROIs) corresponding to individual neuronal cell bodies were isolated from time series using CaImAn (Giovannucci et al 2019). For individual ROIs, the calcium signal per frame was calculated as an average of all pixels within the ROI. This calculation was repeated for each frame in the time series to generate a single-ROI time-course. When more than one ROI per FOV was defined, the same procedure was performed separately for each ROI, resulting in a family of ROI-specific time-courses.

Temporal correlation between eMUA, frequency band power, and individual calcium signals was estimated following a modified procedure from (Watson et al 2018). First, signals (calcium, eMUA, band power) were interpolated to a common time base (fs = 4000 Hz), normalized as *y_Nom_*(*t*) = [*y*(*t*)-y_min_]/*std*(*y*) with *y_min_* as median of the lowest 5% of values in the recording, and segmented into 0.5-s bins. The signal amplitude within the 0.5-s bin was estimated as mean signal amplitude within the bin. Bins were assigned to periods with or without movement and with or without stimulus (0.5 s before and 1.5 s after the stimulation period), where applicable. Correlation between two pairs of signals – on the level of individual bins – was analyzed under the assumption of linear correlation using the ‘corr’ from MATLAB. Pearson’s linear correlation coefficients with a corresponding p-value smaller than 10^-3^ were considered significant.

### Extracellular potential modeling

All neural simulations were done through LFPy 2.3 (Hagen et al 2018) running on NEURON 8.1 (Carnevale & Hines 2006). Mouse cortical L2/3 and L5 pyramidal cell (PC) models were downloaded from the Allen Brain Atlas (https://celltypes.brain-map.org/). The cell models were unmodified, and we used a mixture of models with active conductances throughout the morphology (“all active”), as well as those where the active conductances were present only in the perisomatic region (“perisomatic”). In total, we used 14 different cell models, 5 from L2/3 and 9 from L5.

Action potentials were evoked by a somatic step current injection (POINT_PROCESS in NEURON), where the amplitude of the current was adjusted for each cell model to evoke between two and ten action potentials within 120 ms. Afterward, the transmembrane currents from a 10-ms window around the last spike were extracted, allowing for calculations of the extracellular potential due to the spike at any arbitrary location around the neuron.

Calculations of the extracellular potential were done with LFPy, assuming an electrode diameter of 20 μm, incorporated through the disc-electrode approximation. In experiments, the recording electrodes were embedded in a non-conducting substrate at the cortical surface, which was accounted for in our simulation through the method of images, by multiplying the extracellular potential by a factor of two (Ness et al 2015). The extracellular conductivity was set to 0.3 S/m, and the time step was always 0.0078 ms.

In single-cell simulations (**Fig. 6A**), the neurons were positioned such that the uppermost tip of the apical dendrite was 10 μm below the cortical surface, and the soma was located directly below the recording electrode at the cortical surface.

For the simulated MUA from populations of L2/3 or L5 PCs (**Fig. 6B-C**), we used populations of 1540 cells. We assumed a cell density of 100,000 cells/mm^3^ (Keller et al 2018), which in combination with an assumed layer thickness of 250 μm for both L2/3 and L5 (DeFelipe et al 2002), resulted in a population radius of 140 μm. The 1540 neuronal cell bodies were distributed with a uniform probability distribution within the plane of this cylinder. Since PCs typically extend upwards almost to the cortical surface, the cells were uniformly distributed along the depth axis so that the uppermost tip of the apical dendrite was between 10 and 30 μm below the cortical surface.

The MUA signal was calculated from these populations based on the transmembrane currents extracted from the single-cell simulations described above, and each single-cell contribution was found by convolving the calculated extracellular action potential (EAP) from a single spike with that cell’s spike train. Single-cell spike trains were modeled through Elephant (Denker et al 2018) as independent homogeneous Poisson processes with a refractory period of 5 ms and varying firing rates, and a total duration of 1 s. All 1540 single-cell contributions to the extracellular potential were summed before the resulting extracellular potential was high-pass filtered (4th order Butterworth filter in “filtfilt” mode) above 350 Hz to produce the MUA signal. Finally, the MUA amplitude was calculated as the peak-to-peak amplitude of the simulated MUA signal for each firing rate.

Parameters for the passive and active conductances for the unmyelinated axon model were extracted from the unmyelinated sections of the neuron model from (Hallermann et al 2012) (available from http://modeldb.yale.edu/144526).

The axon model was given a uniform diameter of 0.3 μm, based on observed diameters for intracortical unmyelinated axons (Call & Bergles 2021). Axon models were constructed from sections with lengths of 20 μm each. In all cases, the axon was built of 40 straight, unbranching, and upwards-pointing sections, before a given number of branch points were added at the upper end of the axon model. At each branch point, the two child branches were always pointing in opposite directions in the plane orthogonal to the cortical surface with an angle of 90 degrees between them, forming an approximate Y-shape which was randomly rotated around the cortical depth axis.

Action potentials were evoked in the axon models through a step current injection (POINT_PROCESS in NEURON) to the lowermost part of the axon. The amplitude was −0.25 nA with a pulse width of 0.1 ms, and a 2-ms window around the spike was extracted.

In the single axon example (**Fig. 6D**), the top of the axon was 10 μm below the cortical surface. For the axon population, we used a population radius of 140 μm, the same as for the PC populations. We were unable to find estimates of the density of upwards-pointing afferent axons in L1 and their branching patterns. We therefore assumed an axon density of 0.1 afferent axons per μm^2^: The maximum number of branching points used in this study was four, meaning that each upwards pointing axon had maximally 16 terminal segments. For an axon diameter of 0.3 μm, an axon density of 0.1 afferent axons per μm2 corresponds to such afferent axons at maximum occupying about 50% of the volume in L1 (0.1 μm^-2^ *π *(0.3 μm)^2^ *16 = 0.45), which was regarded to be a reasonable upper bound. This resulted in 6157 afferent axons. Calculation of MUA amplitudes from the axon population otherwise followed the same procedure as for the PC populations.

### Data availability

MATLAB scripts for data analysis and datasets supporting the conclusions of this paper are available from the corresponding authors upon reasonable request. All code to reproduce the simulation results is available from https://github.com/torbjone/cortical_surface_EAPs.

## Results

### Validation of microelectrode array performance in anesthetized mice

Our first goal was to realize wearable PEDOT:PSS surface microelectrode arrays on transparent parylene C substrate for seamless integration with two-photon optical imaging and validate their performance in awake mice with chronic implants. We started by evaluating optical transparency and recording performance of our devices in acute experiments under anesthesia, where experimental conditions were well-controlled. We placed a 32-channel electrode array onto the cortical surface and covered the array with a glass coverslip (see **Methods**; **Fig. 1A-B**). **Figure 1C-D** illustrates two-photon imaging in a mouse expressing EGFP in GABAergic cortical neurons (Tamamaki et al 2003). In addition, cortical astrocytes were labeled with red fluorescent Sulforhodamine 101 (SR101) (Nimmerjahn et al 2004), and Alexa Fluor 680 conjugated to high molecular weight dextran was used as an intravascular contrast agent (Li et al 2020). Under these conditions, we performed two-photon imaging throughout the cortical depth. Although PEDOT:PSS electrode pads and metal leads were not transparent, their shadows gradually disappeared with depth (**Fig. 1D** and **Suppl. Fig. 1**). Alexa Fluor 680, which was excited at 1240 nm, was visible throughout the volume. EGFP, excited at 920 nm, was visible down to ~500 μm due to higher scattering of shorter wavelengths in tissue. The astrocytic marker SR101 was applied to the cortical surface and diffused within astrocytic membranes. It was visible down to ~400 μm due to decreasing labeling intensity with depth. Overall, these results, obtained in the presence of a PEDOT:PSS electrode array on the brain surface, are comparable with the best practice of two-photon imaging of these fluorophores in the mouse cerebral cortex *in vivo* (Kobat et al 2009, Li et al 2019, Nizar et al 2013, Uhlirova et al 2016b).

**Figure 1.**
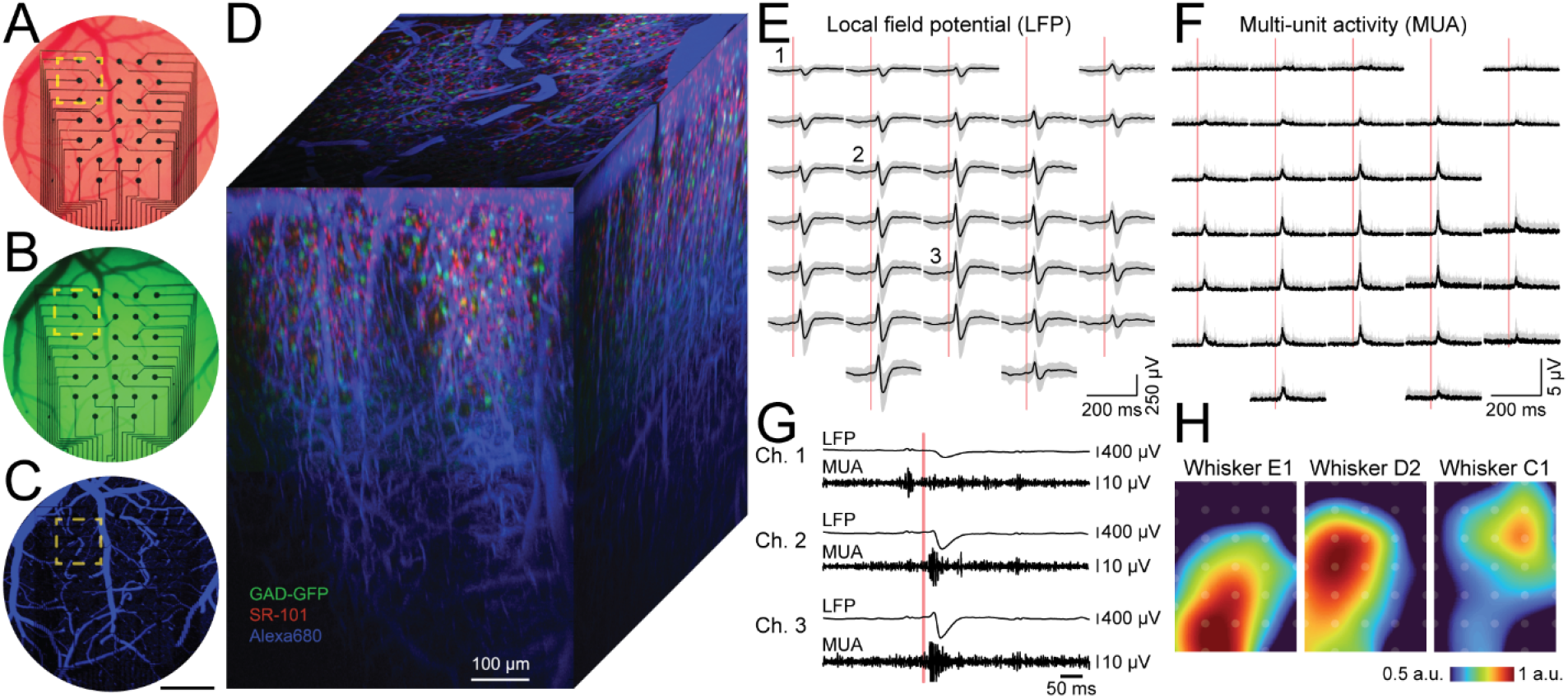
Performance of PEDOT:PSS microelectrode arrays in acute experiments under anesthesia. **A.** Image of the cranial exposure after placement of microelectrode array and glass coverslip. **B.** Epifluorescence image of green fluorescent protein (GFP) in inhibitory (GAD-expressing) interneurons (GAD-GFP). **C.** Maximum intensity projection (MIP) of a low-magnification two-photon image stack of the entire exposure. Blue, intravascular Alexa 680-Dextran. The yellow box outlines the field of view (FOV) shown in panel D. Scale bar, 500 μm. **D.** Three-dimensional reconstruction of a high magnification two-photon image stack, green GAD-GFP; red, Sulforhodamine 101 (SR-101) that labels astrocytes; blue, intravascular Alexa 680 Dextran. Scale bar, 100 μm. **E.** Spatial map of low-pass filtered data (<250 Hz, LFP). The response to stimulation of whisker E1 (vertical red line) is shown. Black lines indicate the mean, grey areas the standard derivation over 50 trials. Traces from two non-working channels (impedance > 5 MΩ) have been removed. **F.** Spatial map of high-pass filtered and rectified data (>350 Hz, MUA). The response to stimulation of whisker E1 (vertical red line) is shown. Black lines indicate the mean, grey areas the standard derivation over 50 trials. Traces from two non-working channels have been removed. **G.** LFP and MUA of a single trial for three channels (indicated 1-3 in (E)) with increasing response amplitude towards the center of the responsive area. **H.** Spatial map of min-max normalized LFP amplitudes in response to stimulation of whisker E1, D2, and C1. The center of the response moves in accordance with the cortical representation of stimulated whiskers.

Next, we evaluated our ability to record LFP and MUA under anesthesia (**Fig. 1E-H**). **Figure 1E-F** illustrates an example of trial-averaged LFP (**Fig. 1E**) and MUA (**Fig. 1F**) responses to deflection of the E1 whisker (n=50 trials). Individual traces corresponding to different recording electrodes are arranged in the figure according to the geometrical layout of the array. Single-trial responses for three out of 30 working channels are shown in **Figure 1G**. Stimulation of different whiskers resulted in movement of the center of mass of the neuronal response in the cortical surface plane (**Fig. 1H**). This is consistent with the well-known whisker representation map within the whisker-barrel cortex (Feldmeyer et al 2013).

Taken together, these data show that PEDOT:PSS surface microelectrode arrays recorded LFP and MUA with high fidelity and offered sufficient optical transparency for deep two-photon imaging.

### Chronic implantation of Wind*an*see devices allow longitudinal measurements in awake mice

To enable longitudinal data acquisition in awake mice, we developed a mechanical solution for installation, connectorization and protection of PEDOT:PSS/parylene C surface electrode arrays aiming for a lifetime of several months while worn by a mouse. Our main consideration was to allow an unhindered access for wide (high numerical aperture) microscope objectives while not obstructing the mouse face including whisker pads.

All parts were designed using CAD (Fusion 360, Autodesk). The shape of the headpost followed a design used for “crystal skull” preparations (Kim et al 2016). The headpost had a notch for installation of a reference screw over the cerebellum as well as three 2-64 UNF-threaded holes (**Fig. 2A** and **Suppl. Fig. 2**). We manufactured the headpost from titanium with a thickness of 1 mm. During implantation surgery, a custom-made installation aid (**Suppl. Fig. 2**) was attached to a stereotaxic arm to allow reproducible placement of the headpost relative to the skull. Parts that secure microelectrode arrays and connector were produced using stereolithography (Form 3, Formlabs). During head fixation, the headpost was attached to the headpost holder (**Fig. 2B** and **Suppl. Fig. 2**). The holder was manufactured from stainless steel and aluminum and consisted of two parts: the bottom part was installed on four Ø1/2-inch optical posts, and the top part was fastened with two 3-56 UNF machine screws to the bottom part.

**Figure 2.**
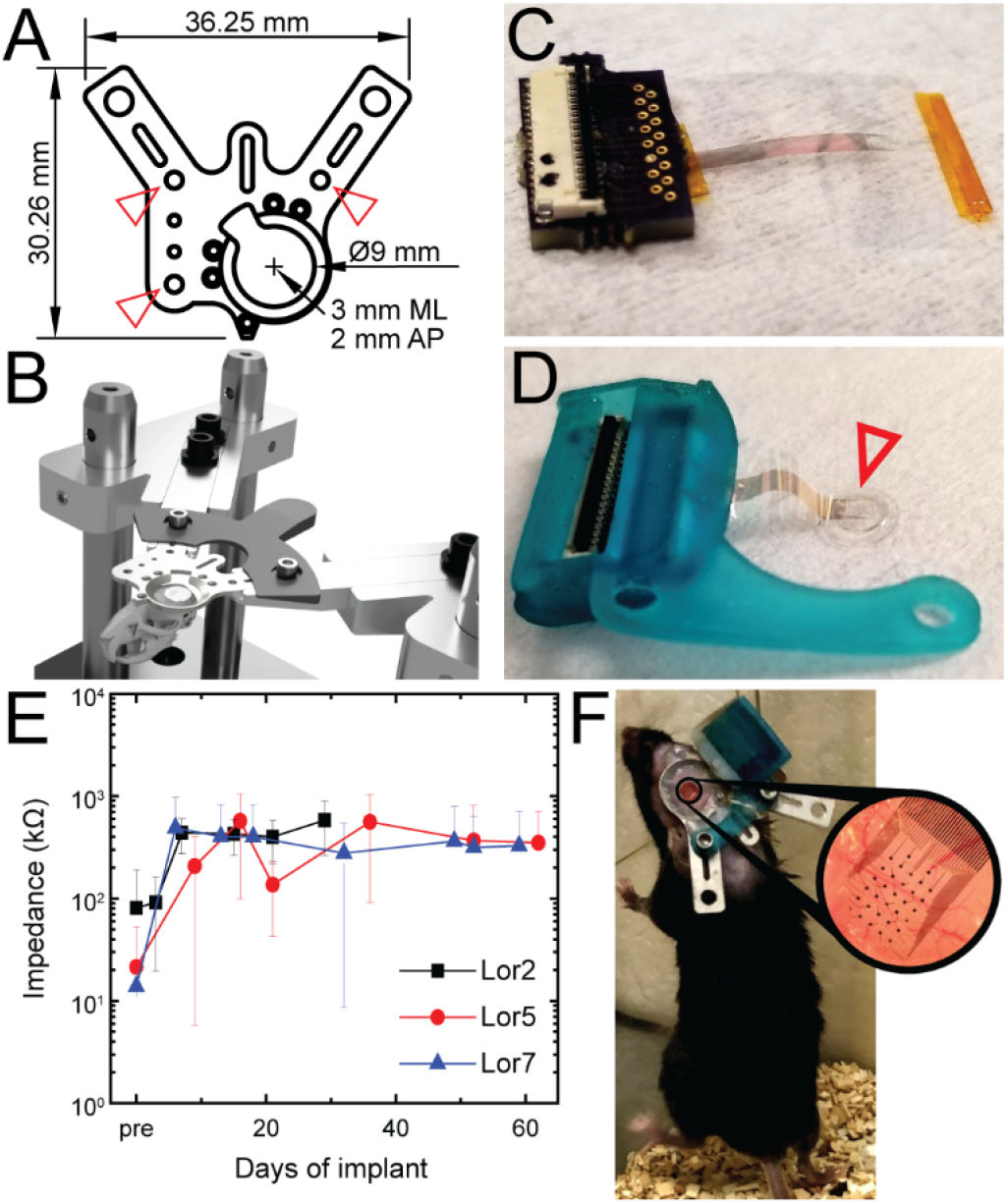
Wearable Wind*an*see devices for chronic implantation. **A.** Schematic drawing of the headpost. The center of the opening is located over the barrel cortex (3 mm mediolateral, 2 mm posterior to Bregma). Threaded holes (red arrowheads) allow fixation of the protective enclosure around the connector board (attached to the Wind*an*see device) to the headpost. The headpost is machined from titanium with a thickness of 1 mm. **B.** Rendering of the stage design. Fixation aid is used to tighten the headpost to the stage. The stage can be fixed to four Ø1/2-inch optical posts. Placing the stage on a goniometer improves alignment of the cortical surface with the horizontal plane during two-photon imaging (not shown). **C.** Photograph of a PEDOT:PSS/parylene C microelectrode array bonded to a connector board with FFC connector. **D.** Photograph of the assembled Wind*an*see device. The microelectrode array has been glued to the glass window (red arrowhead). The connector board has been placed inside a protective 3D-printed case. **E.** Electrode impedance measurements for up to 60 days after implantation. Data for three mice (Lor2, Lor5, Lor7) are shown; data points represent the average impedance of working channels; error bars show the standard deviation. **F.** Photograph of a mouse with implanted Wind*an*see device in its home cage. The animal can move freely when not under head fixation during data acquisition.

The microelectrode array was fused with the bottom 3-mm glass of a glass “window” consisting of two 3-mm and one 5-mm coverslip glass (Goldey et al 2014, Kilic et al 2020) using UV-curable optical adhesive with a refractive index matching that of the glass (Norland Optical Adhesive 61). We coined our wearable devices integrated with the cranial window Wind*an*see.

The array connector (the metal leads on parylene C) was permanently bonded to a small PCB (**Fig. 2C**), and the PCB was encapsulated in a 3D-printed housing (blue in **Fig. 2D, see also Suppl. Fig. 2**). During implantation, the device was permanently attached to the headpost. We observed an increase in impedance of the recording electrodes from below 100 kΩ to ~0.5 MΩ within the first week after the implantation with no significant change afterwards (**Fig. 2E**). Mice carrying implanted devices exhibited normal posture and motor behavior and had no apparent weight loss or other signs of distress under standard housing conditions (**Fig. 2F**).

Overall, wearable Wind*an*see devices and the associated headpost assembly were well tolerated by the mice and remained intact and functional for at least two months.

### Electrophysiological recordings in awake mice with chronically implanted Windansee devices

During recording sessions, awake mice (i.e., no anesthesia or sedation) were head-fixed in a suspended bed, and a ribbon cable was used to connect the Wind*an*see device to the recording equipment (**Fig. 3A**). Head-fixed mice were free to move their body and whisk; we often observed body movement and whisking, spontaneously and in response to the air puff stimulus. Movement of the mouse’s body introduced high-frequency interference in the recorded electrophysiological signal corrupting the MUA band (**Suppl. Fig. 3**). To reduce this motion artifact, we performed post hoc independent component analysis (ICA). For this analysis, we used the raw electrophysiological signal from all 32 recording channels in the array (see **Methods**). For each acquired 180-s run, we calculated the independent components (ICs) and removed up to three IC that that had unrealistically high amplitude while lacking spatial heterogeneity or followed the time-course of laser scanning (**Suppl. Fig. 3**).

**Figure 3.**
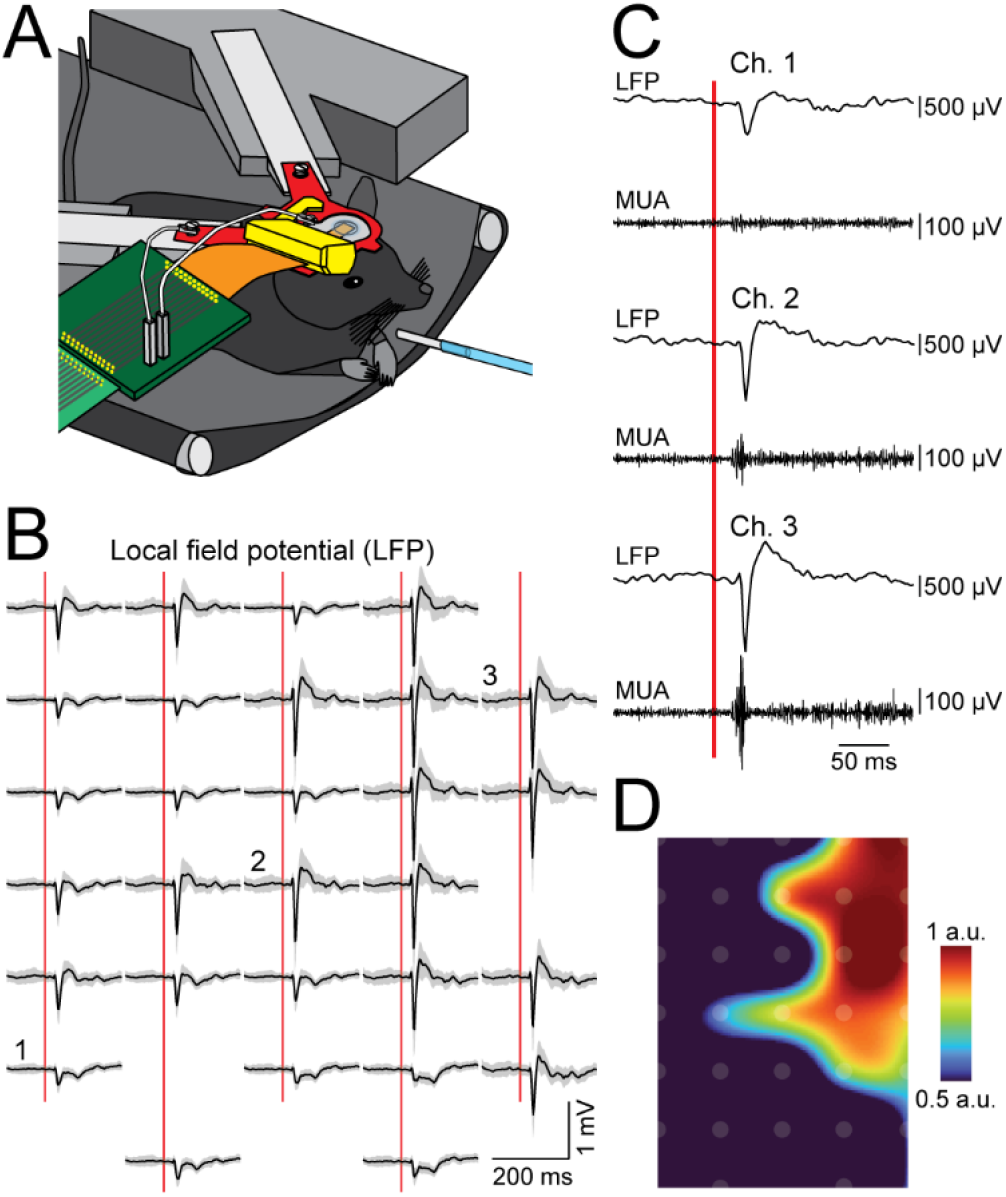
Stimulus-induced MUA and LFP in awake mice with chronically implanted Wind*an*see devices. **A.** Schematic drawing of the recording setup. The animal with implanted Wind*an*see device rests on a suspended hammock while head-fixed to the stage (grey) via titanium headpost (red). The animal can reposition its body and move its limbs. A ribbon cable (orange) couples the connector board inside the protective enclosure (yellow) with the PCB (front left, dark green); the PCB is connected to the amplifier chip (front left, light green). A micro screw inserted into the skull above the cerebellum serves as the reference input. The stage is grounded, and the ground input is connected to the stage. **B.** Spatial map of LFP in response to whisker stimulation with air puff (vertical red line). Black lines indicate the mean, the standard derivation over 30 trials is shown in gray. Traces from three non-working channels have been removed. **C.** Single-trial LFP and MUA responses for three channels indicated in (B). Note increasing response amplitude towards the center of the responsive area. **D.** Spatial map of trial-averaged and min-max normalized LFP amplitudes in response to air puff stimulation.

**Figure 3B-D** shows an example trial-averaged LFP response to air puff stimulation. In **Figure 3B,** individual traces, which correspond to different recording electrodes, are arranged according to the geometrical layout of the array. **Figure 3C** shows low- and high-pass filtered data (i.e., LFP and MUA, respectively) for one individual trial for three channels in the array. Like the results obtained under anesthesia (**Fig. 1**), we observed that the amplitude of the response varied smoothly in space (**Fig. 3D**). The map of the response in **Figure 3D** is more complex compared to that in **Figure 1H.** This is expected, however, from the nature of the stimulus: multiple whiskers sensing the air puff in awake animals vs. well-controlled single-whisker deflection under anesthesia.

Previous studies reported that spiking activity in the cerebral cortex was more correlated with the γ-band (30-80 Hz) compared to α (8-12 Hz) and β (12-30 Hz) bands within the LFP or electroencephalogram (EEG) power spectrum (Burns et al 2010, Watson et al 2018, Whittingstall & Logothetis 2009). Therefore, as an additional validation of the wearable Wind*an*see technology, we examined this correlation in our data (**Fig. 4** and **Suppl. Fig. 4**). We computed LFP spectrograms in the frequency range of 0.05-180 Hz (**Fig. 4A**, top) during spontaneous activity (no stimulus) and extracted time-courses of α-, β-, and γ-band power (**Fig. 4B**). Then, we calculated the correlation (**Fig. 4C**) of each of these time-courses with the MUA envelope (eMUA) that was computed by temporal smoothing of rectified MUA time-course (top trace in **Fig. 4B**, see **Methods**). Consistent with previous reports, the eMUA was stronger correlated with the γ-power compared to α- and β-power: across 33 acquisition runs in three animals, we found significant (p<0.001) correlation between eMUA and α-, β-, and γ-band power in one (~3%), six (~18%), and 16 (~49%) cases, respectively (**Fig. 4D**). In parallel with electrophysiological recordings, we monitored mouse movement using a CCD camera (see **Methods**). We observed that movement (denoted by red bars in **Fig. 4**) typically led to a shift towards higher frequencies in the LFP spectrogram (**Fig. 4A**) coupled to increased MUA (**Fig. 4C**).

**Figure 4.**
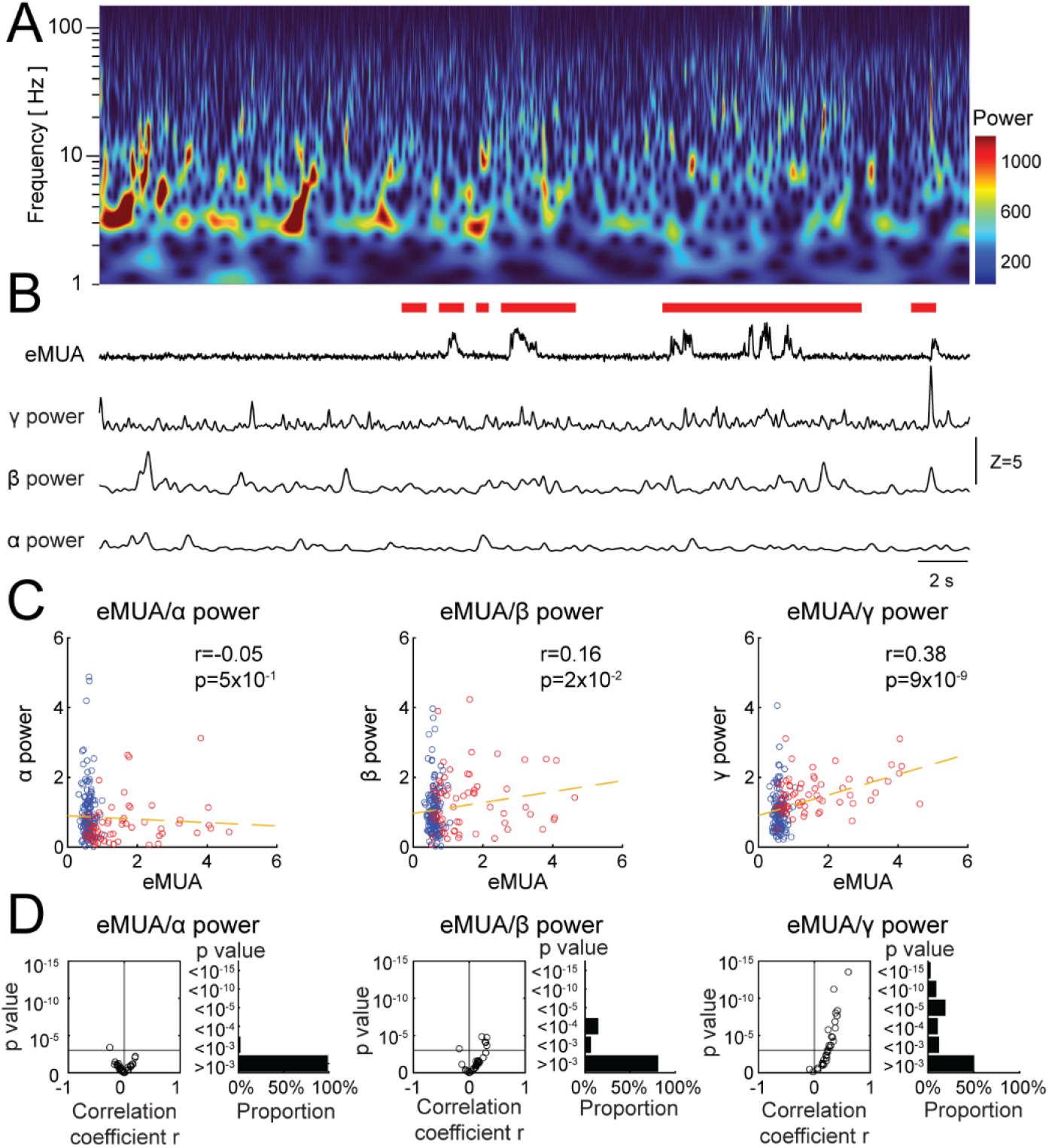
Correlation of MUA with α-, β-, and γ-band of LFP power spectrum. **A.** Spectral LFP composition (1-180 Hz) of a representative 30-s period. **B.** Amplitude of MUA envelope (eMUA) and power of α (8-12 Hz), β (12-30 Hz) and γ (30-80 Hz) bands for the same 30-s period shown in (A). Red bars indicate the animal movement detected with a behavior camera directed towards the animal’s face. **C.** Correlation of eMUA amplitude with power in the α-, β-, and γ-band for the entire 180-s acquisition period (“run”) of which a 30-s excerpt is shown in (A) and (B). For correlation analysis, we calculated the mean within consecutive 0.5-s time bins. Each data point represents an individual 0.5-s bin. Time bins with and without animal movement are shown in red and blue, respectively. The correlation coefficients r and corresponding p-value were estimated for the entire 180-s run. **D.** Summary across 33 runs acquired in three animals: correlation between eMUA and power of α-, β-, and γ-band (left, middle and right, respectively). For each band, a scatter plot of p-value as a function of r is shown on the left, and the p-value distribution on the right. Horizontal lines in the scatter plots indicate the significance threshold (p=10^-3^).

To summarize, wearable Wind*an*see devices offered meaningful MUA and LFP measurements in awake head-fixed mice producing results consistent with existing neuroscience knowledge.

### Simultaneous electrophysiological recordings and two-photon calcium imaging in mice with chronically implanted Wind*an*see devices

Next, we combined electrophysiological recordings with two-photon calcium imaging to address the signal source of MUA recorded at the cortical surface. To enable calcium imaging in neurons, we induced pan-neuronal expression of the calcium biosensor jRGECO1a via local AAV delivery of the hSyn-jRGECO1a transgene into L2/3 of the whisker-barrel cortex. During data acquisition, awake mice were head-fixed under the microscope objective (**Fig. 5A**). MUA, LFP, and calcium imaging data were acquired simultaneously. Because the spike amplitude rapidly decays with distance (Pettersen & Einevoll 2008), we focused on calcium imaging in neuronal cell bodies located in cortical L2/3, 80-250 (151±66) μm below the surface. Single-cell calcium time-courses were extracted from ~130×110 μm FOVs (see **Methods**). For quantification of correlation between MUA and single-neuron calcium activity, we computed eMUA as in **Figure 4**. All neurons in the same FOV were assigned to the same corresponding eMUA recorded from the nearest surface electrode above the imaging plane (**Fig. 5B**).

**Figure 5.**
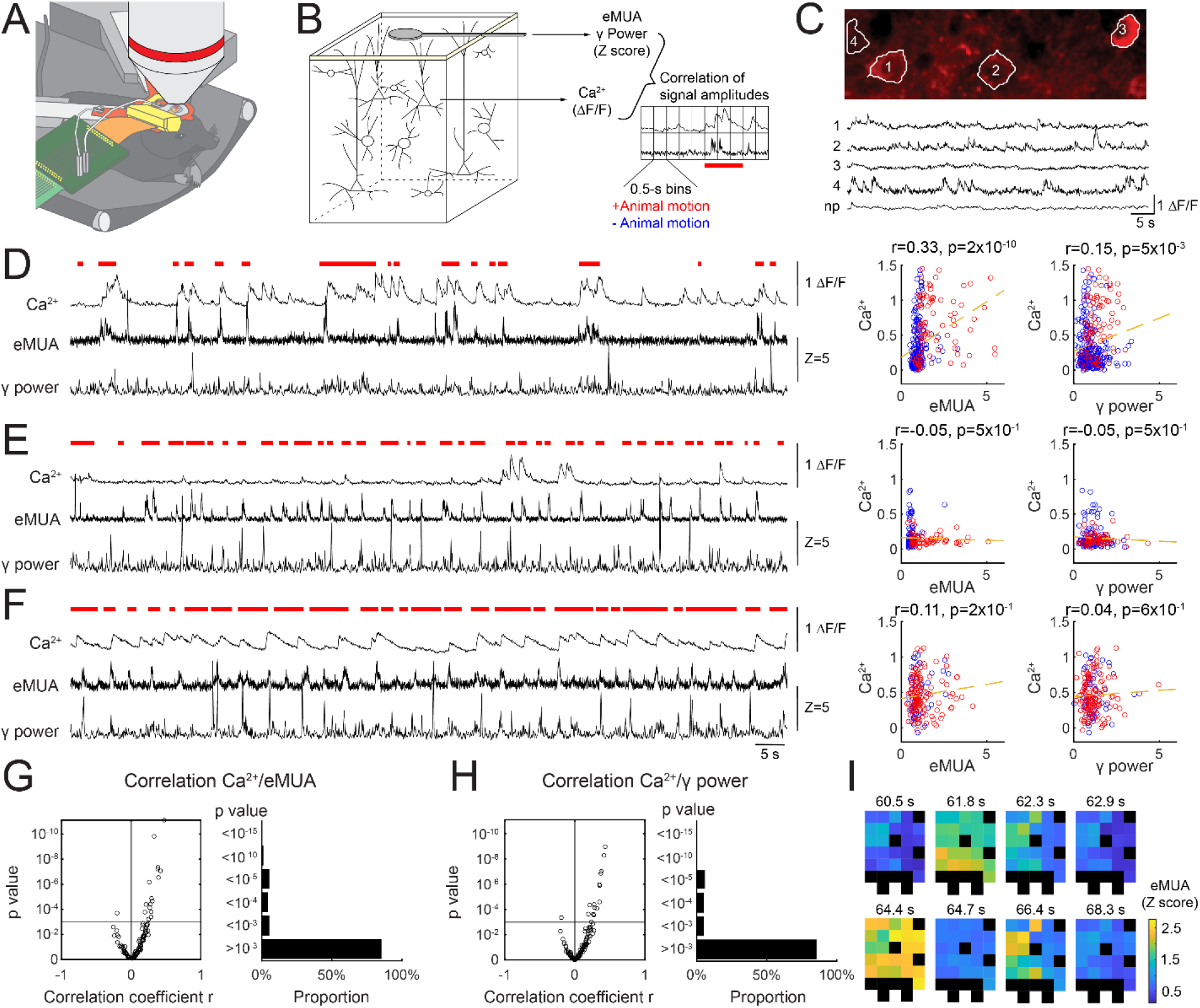
Correlation between MUA and calcium activity of individual L2/3 neurons. **A.** Schematic drawing of the experimental setup. Stage and recording equipment were placed inside the two-photon microscope enclosure for simultaneous calcium imaging and electrophysiological recordings. **B.** Schematic drawing of the experimental approach. Calcium signals from jRGECO1a-expressing neuronal cell bodies were correlated with electrophysiological signals simultaneously recorded by the nearest surface microelectrode above the imaging plane. For correlation analysis, time-courses were segmented into 0.5-s time bins and assigned to periods with (red) or without (blue) animal movement. **C.** Top: a representative imaging FOV with several outlined ROIs. This image was computed as an average of the acquired image time series; the contrast is due to jRGECO1a fluorescence. Bottom: time-courses of calcium activity extracted from the ROIs outlined in the image on top and neuropil (np, corresponding to all pixels outside the ROIs). **D-F**. Left: three examples of eMUA, γ-power, and single-neuron calcium time-course. Red bars indicate animal movement. Right: correlation of eMUA or γ-power with calcium signal in 0.5-s bins assigned to periods with (red) and without (blue) movement corresponding to the cases shown on the left. For each case, the correlation coefficients r and corresponding p-value were estimated for the entire 180-s run. **G.** Summary plots for correlation between single-neuron calcium activity and eMUA: a scatter plot of p-value as a function of r (left), and the p-value distribution (right) for 136 sampled neurons (somatic ROIs) from 33 runs in three animals. The horizontal line on the scatter plot indicates the significance threshold (p=10^-3^). The correlation is significant (p<10^-3^) for 18 of 136 (~13%) sampled neurons. **H.** Same as in (G) for correlation between single-neuron calcium activity and γ-power. The correlation is significant (p<10^-3^) for 18 of 136 (~13%) sampled neurons. **I.** Spatiotemporal dynamics of eMUA amplitude (shown as Z score) across the microelectrode array. Non-working channels are replaced with black tiles.

Calcium time-courses extracted from regions of interest (ROIs) corresponding to individual neuronal cell bodies exhibited variability (**Fig. 5C**), consistent with variability in spiking across neurons observed in electrophysiological studies (Watson et al 2018). Surprisingly, only a small fraction of neurons faithfully followed eMUA (**Fig. 5** and **Suppl. Fig. 5**). Out of 136 segmented neuronal cell body ROIs across 33 recordings in three mice (3.4±1.5 segmented neuronal cell body ROIs per FOV), only 18 (~13%) showed significant (p<10^-3^) correlation with eMUA. One of these neurons is shown in **Figure 5D**. To compute correlation, we divided the time-course into consecutive 0.5-s time bins. Each point in the scatter plot in **Figure 5D** (right) shows a correlation value for one such time bin. Like in **Figure 4**, we monitored mouse movement (see **Methods**) and classified time bins into those with and without movement (red and blue points, respectively, in the scatter plot).

Most sampled neurons showed no significant correlation with eMUA, irrespective of their level of activity or presence/absence of mouse movement. Example neurons with low and high activity levels but no significant correlation with eMUA are shown in **Figure 5E** and **Figure 5F**, respectively. Correlation statistics for the entire sample of 136 segmented cell body ROIs is shown in **Figure 5G**, where the correlation coefficient *r* is plotted against the p-value. Most of the observations (118 out of 136; 87%) have *p*>10^-3^, indicating non-significant correlations. A similar distribution was found for the correlation between Ca^2+^ signals and γ band power (**Fig. 5H**). While in the present study we focused on a single electrode above the imaging FOVs, the presence of multiple electrodes in the array would allow extending this analysis across the cortical columns and regions (**Fig. 5I**).

### MUA recorded at the cortical surface is likely to reflect spikes in L1 afferents

Given the low probability of finding L2/3 neurons that were significantly correlated with MUA recorded at the surface (**Fig. 5**), we turned to computational modeling to evaluate alternative candidate biophysical processes that could account for the MUA signal: spiking of local cortical neurons and afferent axons.

First, we considered cortical pyramidal cells (PCs). We used models of morphologically reconstructed (biophysically detailed) L2/3 (n=5) and layer 5 (L5, n=9) mouse PCs from the Allen Brain Atlas (https://celltypes.brain-map.org/). These model neurons were endowed with passive and active membrane conductances including those supporting back-propagation of action potentials (see **Methods**). Two example neurons from L2/3 and L5 are shown in **Figure 6A** (top and middle panels). Spiking in the cell models was induced by depolarizing the soma with a step current, and the resulting extracellular action potential (EAP) was calculated at the cortical surface (see **Methods** and **Suppl Fig. 6A**). In general, PCs with somas located deeper below the surface produced smaller surface EAP (**Fig. 6A**, bottom).

**Figure 6.**
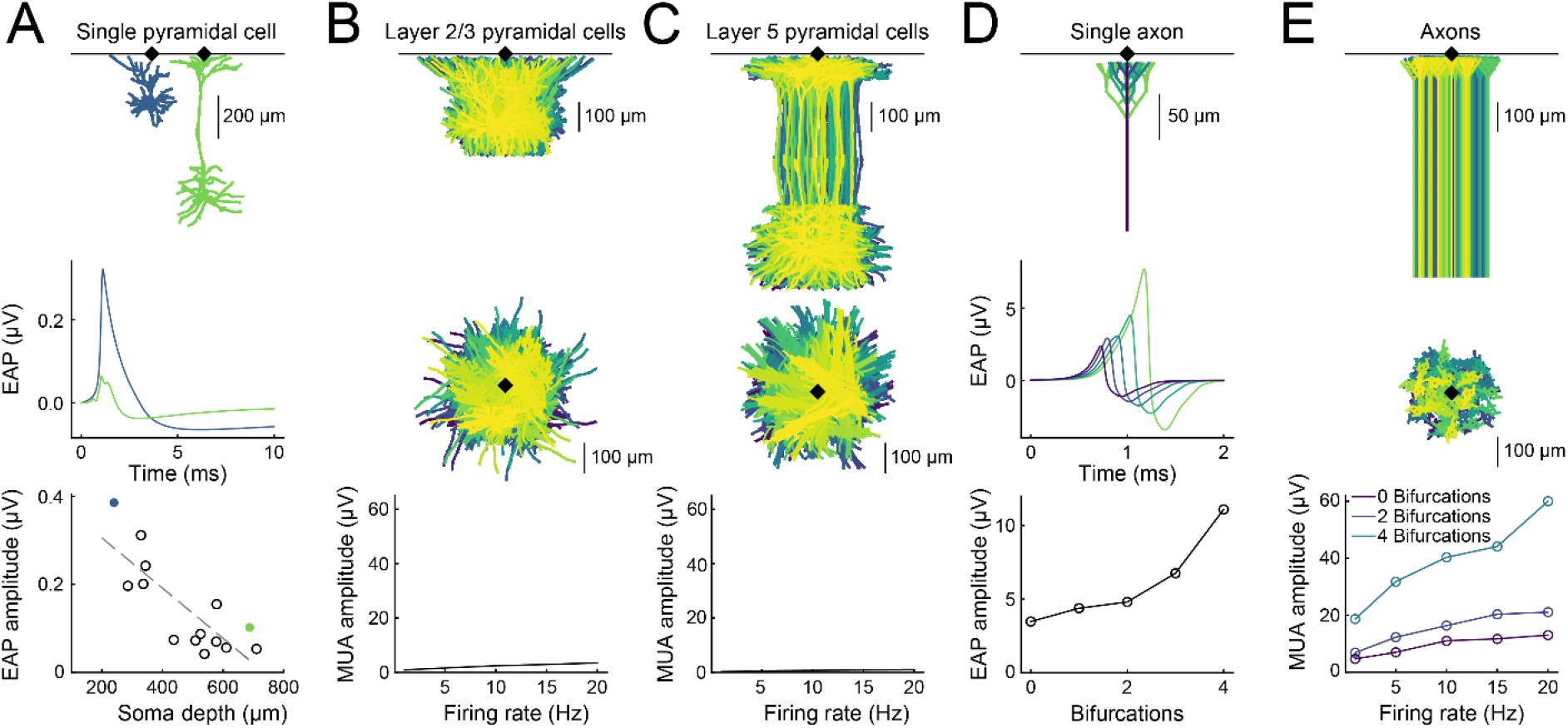
Modeling of EAP and MUA at the cortical surface due to firing of local neurons and afferent axons. **A.** Top: example cell models of PCs from L2/3 (blue) and L5 (green) from the Allen Brain Atlas database. Middle: spiking in the cell models were induced by depolarizing the soma with a step pulse, and the resulting EAP was calculated at the cortical surface. Bottom: the peak-to-peak EAP amplitude at the cortical surface plotted against soma depth for all used cell models. The dashed gray line shows a linear fit. All cell models used here generated EAP amplitudes below 0.5 μV at the cortical surface. **B.** A population of (identical) L2/3 PCs (corresponding to the L2/3 cell shown in (A)) with soma locations confined within a cylinder (radius: 140 μm). One hundred example cells are plotted seen from side (top) and top (middle). The MUA amplitude at the cortical surface, here quantified as the peak-to-peak amplitude of the simulated MUA signal, increased with firing rate (bottom). **C.** Same as in (B), but for a population of (identical) L5 PCs (corresponding to the L5 cell shown in (A)). **D.** A single unmyelinated axon (fixed diameter of 0.3 μm), with different degrees of bifurcation (top). Axonal bifurcations substantially increase the EAP (middle and bottom panel). **E**. A population of 6157 axons within a cylinder (radius: 140 μm). Three hundred axons with different numbers of bifurcations (0, 2 and 4 bifurcations) are shown from the side in (top) and from above (middle). The MUA amplitude at the cortical surface, here quantified as the peak-to-peak amplitude of the simulated signal, increased with the number of bifurcations and with the firing rate (bottom).

Because a surface electrode has a certain “listening area” around it (Buzsaki et al 2012), we simulated a population of 1540 cells with a density of ~10^5^ cells/mm^3^ (Keller et al 2018), with either L2/3 or L5 PCs corresponding to the neurons shown in **Figure 6A** (top) and with somas confined within a column of radius 140 μm in the cortical surface plane (xy) and the uppermost tip of the apical dendrite between 10 and 30 μm below the cortical surface (see **Methods**). For each neuron, spikes were Poisson-distributed with a refractory period of 5 ms with a given mean firing rate. Different neurons were assumed to generate spikes independently of each other. **Figures 6B-C** show results for the L2/3 and L5 populations, respectively (see also **Suppl. Figure 6B**). The MUA amplitude at the cortical surface, quantified as peak-to-peak amplitude of the simulated signal, increased with the mean firing rate (1-20 Hz).

Next, we considered the surface EAP due to spiking of L1 afferents. Cerebral cortex receives both myelinated and unmyelinated afferents. Myelinated axons, which usually enter from subcortical white matter ascending vertically in cortical columns, lose their myelin sheet upon reaching L1 (van Tilborg et al 2017). Therefore, most axonal segments found in L1 are unmyelinated. In **Figure 6D**, we model individual unmyelinated axons while varying the number of branches. Axonal branching substantially increased the surface EAP amplitude (McColgan et al 2017): a single axon with a fixed diameter of 0.3 μm and four bifurcations within the top 50 μm produced an EAP threefold higher compared to an unbranched axon.

Then, we simulated a population of 6157 unmyelinated axons within the same cylinder with a radius of 140 μm (300 axons are shown in the top row of **Fig. 6E**) while varying the number of branches in L1 (0, 2 and 4 bifurcations). The MUA amplitude at the cortical surface, quantified as the peak-to-peak amplitude of the simulated signal, increased with the number of bifurcations and with the mean firing rate of the axons (1-20 Hz) (**Fig. 6E** and **Suppl. Fig. 6C**, left). As for the case of PCs, each axon’s spikes were Poisson distributed with a 5-ms refractory period with a given mean firing rate, and different axons were assumed to generate spikes independently of each other. MUA signals generated by the population of axons were over an order of magnitude larger compared to that generated by the population of L2/3 PCs (compare the bottom plot in **Fig. 6E** to that in **6B**). Some cortical L1 afferents may retain their myelin sheet until the axonal terminals. Therefore, we also tested myelinated axons, which gave similar amplitudes as unmyelinated axons (**Suppl. Fig. 6C**).

Another potential source of the surface MUA signal are inhibitory neurons (INs) with cell bodies located in L1. Because of their proximity to the surface, EAP due to a spike in a L1 neuron can be much larger than that from L2/3 or L5 PCs (**Suppl. Fig. 6D**). However, density of L1 INs is very low, ~60 cells within a cortical column with a radius of ~140 μm (Meyer et al 2010). Assuming a recording electrode with a listening sphere radius of ~100 μm, spikes in ~15 L1 INs would be detected. In practice, the EAP amplitude decreases with distance ***r*** from the electrode scaling in-between ***1/r*** and ***1/r^2^*** (Pettersen & Einevoll 2008). Therefore, even fewer L1 INs would produce detectable signals. This back-of-envelope calculation suggests that if L1 INs could be observed, it would be as single units. However, we were unable to reliably isolate units from our experimental data.

Taken together, our simulation results put forward spiking of L1 afferents as the most likely biophysical process underlying MUA recorded at the cortical surface by Wind*an*see electrodes.

## Discussion

### Summary

The recent advent of neurophotonics (Abdelfattah et al 2022) and high-yield electrophysiological recordings (Vazquez-Guardado et al 2020) has brought into sharp focus the question of sampling biases specific to each measurement modality. These biases confound optical and electrophysiological experimental results complicating the interpretation of data across studies (Siegle et al 2021). Transparent electrode arrays, such as our Wind*an*see technology, are uniquely suited to reconcile optical and electrophysiological measurements of neuronal activity. In the present study, we explored the etiology of MUA detected at the cortical surface *in vivo* in fully awake mice with Wind*an*see devices implanted over the whisker-barrel cortex. We focused on neuronal cell bodies located in L2/3 because their activity is commonly believed to underly spikes picked up by surface electrodes (Khodagholy et al 2015). Combining Wind*an*see recordings with two-photon calcium imaging, we show that only a minority of these neurons faithfully followed MUA recorded above the imaging FOV. Searching for alternative, more likely sources of the MUA signal, we used computational modeling to simulate EAP and MUA due to spiking of different types of neurons as well as L1 afferents (i.e., input axons). Our computational results indicate that spikes in L1 afferents is a more plausible source of surface MUA compared to spikes in L2/3 PCs or other local neurons (L5 PC and L1 INs).

### Development and validation of wearable Windansee devices

The central component of the Wind*an*see device is a surface PEDOT:PSS/parylene C electrode array. The fabrication procedure included a thin Au nanorod layer underneath PEDOT:PSS electrodes that was previously shown to improve the adhesion of the polymer, preventing delamination and increasing longevity of the device (Ganji et al 2018). Following up on our previous demonstration of biocompatibility of these electrode arrays (Ganji et al 2018), in the present study, we show that chronically implanted devices remained functional for months offering minimally invasive longitudinal electrophysiological recordings as well as a see-through cranial window for optical imaging. Developing a lightweight, robust headpost assembly was an important factor contributing to this success. A relatively small size (20-μm diameter) and low impedance (<0.5 MΩ) of PEDOT:PSS electrodes enabled recordings of both stimulus-induced and spontaneous, space-resolved MUA and LFP.

As part of the validation of wearable Wind*an*see technology, we examined spontaneous MUA in the context of the ongoing LFP power spectrum, which is commonly used to describe internal brain states (Lee & Dan 2012). An increase in neuronal firing rate during elevations in γ-oscillations (30-100 Hz) has been reported in prior electrophysiological studies that used penetrating electrodes in order to isolate single units (Nir et al 2007, Watson et al 2018) or record MUA (Burns et al 2010, Whittingstall & Logothetis 2009). In agreement with these reports, we observed stronger correlation of spontaneous MUA with fluctuations in γ-oscillations compared to lower frequencies (8-30 Hz).

### Application of Windansee to investigate etiology of surface MUA

Extracellular electrophysiological recordings cannot be used to quantify silent (or nearly silent) neurons; neurons that do not spike remain invisible for this measurement modality. Two-photon calcium imaging, on the other hand, allows measurements from all neurons in the FOV irrespective of their level of activity. With rapid development and dissemination of new neurophotonic and electrophysiological methods, there is a growing need to understand the relationship between these measurements to enable meta-analysis across measurement modalities (Siegle et al 2021). In the present study, we focused on the cellular origin of spikes detected from the cortical surface leveraging high optical transparency of Wind*an*see devices.

In neurons, somatic calcium signals (i.e., extracted from the cell bodies) are driven by voltage-gated calcium channels that open during the depolarizing phase of an action potential (Grienberger & Konnerth 2012). Therefore, these signals reflect spikes rather than subthreshold events (Vogelstein et al 2010). We examined neuronal somas in L2/3 that have been proposed to generate EAPs detectable from the cortical surface (Khodagholy et al 2015). Contrary to our expectation, only a small percentage of these neurons exhibited spontaneous calcium activity significantly correlated MUA. For most sampled neurons this correlation was low suggesting that other sources may account for the surface MUA signal.

### MUA recorded from the cortical surface is likely to reflect L1 inputs rather than firing of local neurons

Knowing the morphology of a neuron and the membrane ionic properties allows forward calculation of the extracellular potential generated by transmembrane currents. We employed computational modeling to predict EAP at the cortical surface generated by biophysically detailed L2/3 PCs, L5 PC, L1 INs and axons projecting to L1. Then, we calculated the corresponding surface MUA assuming the literature values for cell and axonal density and a broad range of firing rates. For neurons, we leveraged the Allen Brain Atlas library of morphologically reconstructed cell models from the mouse cortex. Unmyelinated axons were modeled as cylinders with a fixed diameter of 0.3 μm ascending vertically to L1 along the cortical axis and bifurcating within the top 50 μm. Using multicompartmental modeling, we first computed the transmembrane currents and then calculated the extracellular potential based on volume conduction theory (Einevoll et al 2013, Holt & Koch 1999).

The EAP amplitude was shown to decrease with distance ***r*** from the electrode scaling in-between ***1/r*** and ***1/r^2^*** (Pettersen & Einevoll 2008). For realistic neurons, however, the fall of the EAP with distance from the soma depends on exact morphology and active membrane conductances supporting dendritic backpropagation of action potentials (Stuart et al 1997). Our computational results show that individual L1 INs as well as individual L1 axons generate surface EAPs substantially larger than that of individual L2/3 or L5 PCs. Due to their low density, L1 INs would be detected as single units rather than MUA. In contrast, L1 axons have high density and firing rates, therefore producing large-amplitude MUA. Taken together, these computational results put forward spiking of L1 axons as the dominant source of the surface MUA signal, although L1 INs can still provide a minor contribution.

These computational results explain the apparent discrepancy between two-photon measurements of L2/3 spiking activity and surface MUA: the MUA signal reflects the coherence of the input rather than that of local neurons, where individual local neurons may or may not fire in response to the input. This scenario is consistent with large variability in spiking across neurons observed in prior electrophysiological studies (Watson et al 2018). Our conclusion of spikes in afferent axons contributing to cortical MUA is also in agreement with an earlier study that used laminar multielectrodes in the whisker-barrel cortex of anesthetized rats (Vinokurova et al 2018). This study reported that some of the MUA response to whisker stimulation remained following pharmacological blockade of cortical glutamatergic receptors indicating sensitivity to afferent spikes.

### Limitations of the present study

The present study has several limitations. The calcium biosensor jRGECOa1 was expressed after local AAV injection under control of the human synapsin promoter resulting in non-selective neuronal expression. In the future, cell-type-specific expression of (color-multiplexed) calcium or voltage probes in mice with chronic Wind*an*see implants will allow addressing the role of genetically identified neuronal cell types to generation of the extracellular potential (Hagen et al 2016, Uhlirova et al 2016a). One additional caveat is that the level of expression of voltage-gated calcium channels varies across neuronal cell types, affecting the signal to noise ratio of the imaging signal (Langer & Helmchen 2012).

Another experimental limitation of the present study was high-frequency interference affecting MUA signals during movement of the mouse. To mitigate this artifact, we decomposed MUA using ICA and removed ICs that were consistent with motion artifacts or interference with imaging instrumentation. Although we were careful to remove only ICs timed to the animal movement only when they lacked spatial heterogeneity, we cannot rule out that this procedure reduced some of the true, movement-induced spiking activity. Optimization of the Wind*an*see device assembly in the future will improve mechanical stability diminishing this artifact. Because the microelectrode arrays were fused with the glass window, electrodes could move along the cortical surface when movement of the mouse body produced motion of the brain inside the skull. This motion can be suppressed by adding to the thickness of the glass insert (Andermann et al 2013), at the price of eliminating the space that is normally present between the brain surface and overlaying cranial bone. Further efforts will be required to find the right compromise between these factors. Tighter adherence of electrodes to the brain surface would also improve our ability to isolate single units that have been detected with NeuroGrid devices (Khodagholy et al 2015).

### Outlook

Combined with two-photon calcium imaging, Wind*an*see technology offers quantification of single-cell activity in the context of space-resolved brain rhythms without insertion of wires into the brain tissue. In the present study, we used 32-channel arrays covering ~ 1 mm^2^ area within the whisker-barrel cortex. Scaling up of Wind*an*see devices to cover multiple functionally distinct cortical areas would enable capturing large-scale phenomena, e.g., entrainment of vasomotion by γ-oscillations that has been hypothesized to underly cortical parcellation in “resting state” hemodynamic studies (Machler et al 2021, Mateo et al 2017). Importantly, surface electrode arrays can be scaled up without increasing the risk of damage to the vascular bed.

This technology offers space-resolved “gold standard” measures of neuronal electrical activity to validate or supplement physiological variables that can be imaged optically using specific synthetic or genetically encoded probes as well as intrinsic contrasts. In the present study, we performed simultaneous electrophysiological recordings and calcium imaging. In the future, Wind*an*see devices will be used in combination with imaging of other optical probes including those of neuromodulatory neurotransmitters (Feng et al 2019, Jing et al 2018, Sabatini & Tian 2020, Sun et al 2018, Wan et al 2020) as well as optical imaging of vascular, metabolic and hemodynamic activity (Machler et al 2021). This multimodal paradigm will help in addressing important questions of neuronal circuits underlying behavior, improve the interpretability of LFP (Buzsaki et al 2012, Einevoll et al 2019, Einevoll et al 2013), underpin noninvasive imaging modalities (Uhlirova et al 2016a), and aid in the development of data-driven computational models (Billeh et al 2020, Einevoll et al 2007, Gagnon et al 2015, Hagen et al 2016).

## Acknowledgements

We gratefully acknowledge support from the National Institutes of Health (BRAIN Initiative R01NS123655 to SAD, BRAIN Initiative U19NS123717 to AD, BRAIN Initiative R01MH111359 to AD, BRAIN Initiative UG3NS123723 to SAD, NIH R01DA050159 to AD, NIH DP2-EB029757 to SAD) and National Science Foundation (1728497 to SAD, 1351980 to SAD). This work was also supported by the European Union Horizon 2020 Research and Innovation Programme under (Grant Agreement No. 945539 to GTE) and Human Brain Project (SGA3 to GTE). We thank Qun Chen, Kimberley Weldy, and Payam Saisan for technical support. We are grateful to Yuchio Yanagawa who generously shared with us transgenic GAD67-GFP mice. The authors acknowledge that some of the data analysis reported in this paper was performed on the Shared Computing Cluster which is administered by Boston University’s Research Computing Services.

