## Supplementary Figures 1-6 for "Imaging through Wind*an*see electrode arrays reveals a small fraction of local neurons following surface MUA"

**Supplementary Figure 1. Quality of two-photon imaging at different depths below the PEDOT:PSS/parylene-C microelectrode array**

Left: high magnification two-photon maximum intensity projection (MIP) of consecutive 150-µm slabs from the surface to a depth of 750 µm for the same data as in **Figure 1A-D**. Green, green fluorescent protein (GFP) in inhibitory (GAD-expressing) interneurons (GAD-GFP); red, Sulforhodamine 101 (SR-101) that labels astrocytes; white, intravascular Alexa 680-Dextran. Locations of the microelectrodes and leads on the surface are indicated by yellow dotted lines. Scale bar, 100 µm. Right: Orthogonal (XY) MIP of intravascular Alexa 680-Dextran acquired from surface (top) to 800 µm below the surface (bottom) corresponding to the region outlined in yellow in **Figure 1A-C**. Scale bar, 100 µm.

**Supplementary Figure 2. Design of essential components of the head fixation system**

**A-B.** Rendering (A) and schematic drawing (B) of headpost with installation aid.

**C-D.** Rendering (C) and schematic drawing (D) of headpost with 3D-printed enclosure for the connector board. The parts shown in (C) and (D) are permanently affixed to the animal's skull. The inset within the red dotted rectangle in panel D is a cross-section of headpost and enclosure with connector board across the red dotted line. Individual components of the assembly are color-coded in the cross-sectional view.

**E-F.** Rendering (E) and schematic drawing (F) of headpost, 3D-printed enclosure, and stage for head fixation and fixation aid. The inset within the red dotted rectangle in panel F is a cross-section of headpost, enclosure, stage, and fixation aid across the red dotted line. Individual components of the assembly are color-coded in the cross-sectional view.

**Supplementary Figure 3. ICA-based denoising of electrophysiological data**

**A.** An example of raw electrophysiological data from one electrode over a period of ten stimulus trials where single air puffs were delivered (red lines indicate the stimulus). Top and bottom show data before and after denoising (through removing three ICs). Yellow shading indicates periods where movement was detected from the surveillance video.

**B.** Top: raw electrophysiological data (left) and isolated LFP (center) and MUA (right) over a 2-s peri-stimulus period (red line). The trial average is shown below ("avg"). Bottom: the same as above after denoising by removal of three ICs. Periods of mouse movement are marked in yellow. Note the different scale of MUA before and after denoising.

**C**. The first 16 ICs (of 32) are shown for a 2-s per-stimulus period for trial 4 (highlighted in blue in (B)). ICs C5 and C13 (blue) (and C26, not shown) were removed from the data.

**D.** Spatial maps of the first 16 ICs. Note the uniform contribution of high-amplitude C5 (likely representing a motion artifact) while C13, which mirrors the time-course of laser scanning, has higher amplitude over the region that was imaged.

**Supplementary Figure 4. Correlation of eMUA with α-, β-, and γ-band power**

Further examples related to **Figure 4**. The left-hand side of A-C shows the LFP spectrogram and time-courses of eMUA α-, β-, and γ-band power over a 30-s period. Red bars indicate the animal movement detected with a surveillance camera. The right-hand side of A-C shows bin-wise correlation between eMUA and α-, β-, and γ-band powers over the entire 180-s acquisition run. Data points represent average signal amplitudes within individual 0.5-s time bins with (red) or without (blue) movement.

**Supplementary Figure 5. Correlation of eMUA and γ-band power with single-neuron calcium activity**

Further examples related to **Figure 5**. The left-hand side of (A)-(C) shows single-neuron time-courses of calcium activity along with electrophysiological recordings (eMUA, and γ-band power) from a nearby surface electrode. The right-hand side of (A)-(C) shows bin-wise correlation of eMUA or γ-band power with single-neuron calcium activity across the entire 180-s run. Data points represent average signal amplitudes within individual 0.5-s time bins with (red) or without (blue) movement. The correlation coefficients r and corresponding p-value were estimated for the entire 180-s run.

**Supplementary Figure 6. Modeling of MUA**

**A.** One single L2/3 PC is shown in the top panel (same cell as in **Figure 6A**). The panel in the middle shows the membrane potential ($V_m$) at the soma (green) and the apical dendrite before it bifurcates (magenta). The bottom panel shows the corresponding EAP measured at the cortical surface (black).

**B**. Example simulated MUA time-courses of L2/3 (left column) or L5 PCs (right column), before (black) and after (red) applying a high-pass filter (>350 Hz, see **Methods**) from a population of identical cell models (N=1540 cells within a population radius of 140 μm). The neuron models are firing at different rates between 1 and 20 Hz (rows). MUA amplitudes in **Figure 6B-C** were calculated as the peak-to-peak amplitude of the high-pass filtered (red) MUA signal for each population at each firing rate.

**C**. Example simulated MUA time-courses from a population of unmyelinated (3 columns on the left) or myelinated (3 columns on the right) axon models (N=6157 axons within a radius of 140 μm), either from before (black) or after (red) applying a high-pass filter (> 350 Hz, see **Methods**). The axon models are firing at rates between 1-20 Hz (rows). For each axon type (unmyelinated or myelinated), we show results from either 0, 2, or 4 bifurcations near the cortical surface (columns).

**D**. Left: a single L1 neuron is shown in the top panel. Spiking was induced by depolarizing the soma with a step pulse, and the resulting EAP was calculated at the cortical surface (bottom panel, left). A population of L1 neurons (N=60 cells within a population radius of 140 μm) seen from the side and top (upper center panel), as well as the simulated MUA amplitude as a function of the firing rate of the simulated neurons (lower center panel). The MUA amplitude is calculated as the peak-to-peak amplitude of the simulated time course of the high-pass filtered (>350 Hz, see **Methods**) MUA signal (right column, red lines) at firing rates between 1 and 20 Hz (rows).

### Supplementary Figure 1

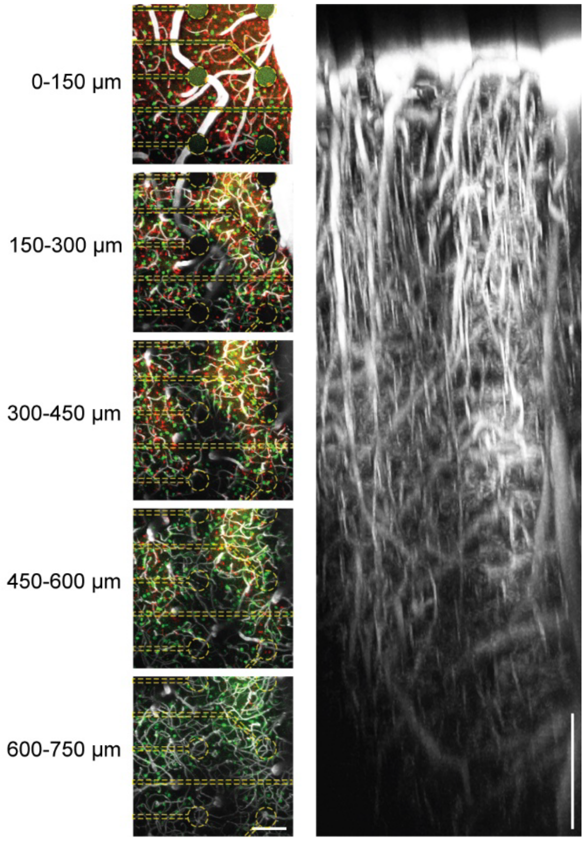

### Supplementary Figure 2

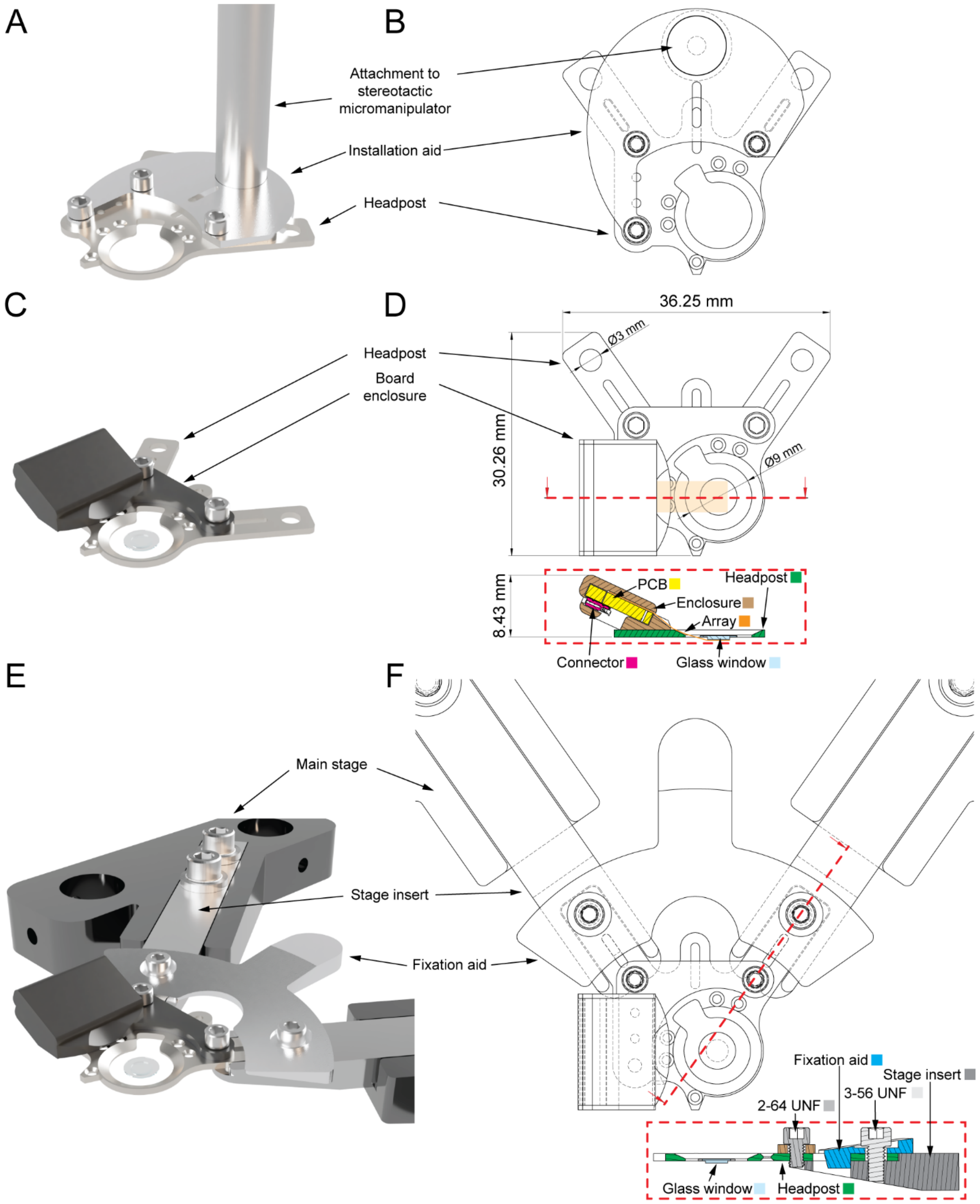

### Supplementary Figure 3

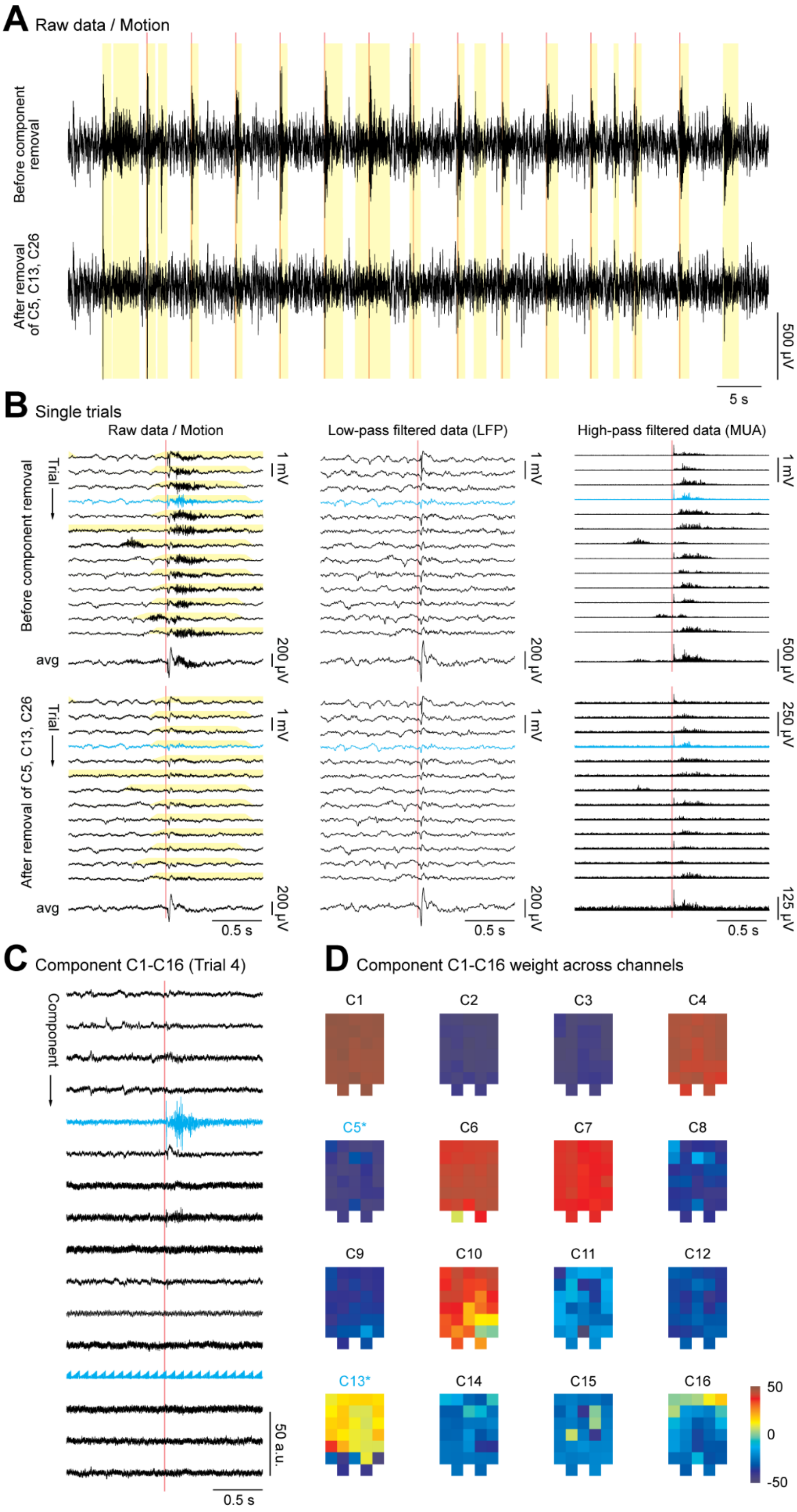

### Supplementary Figure 4

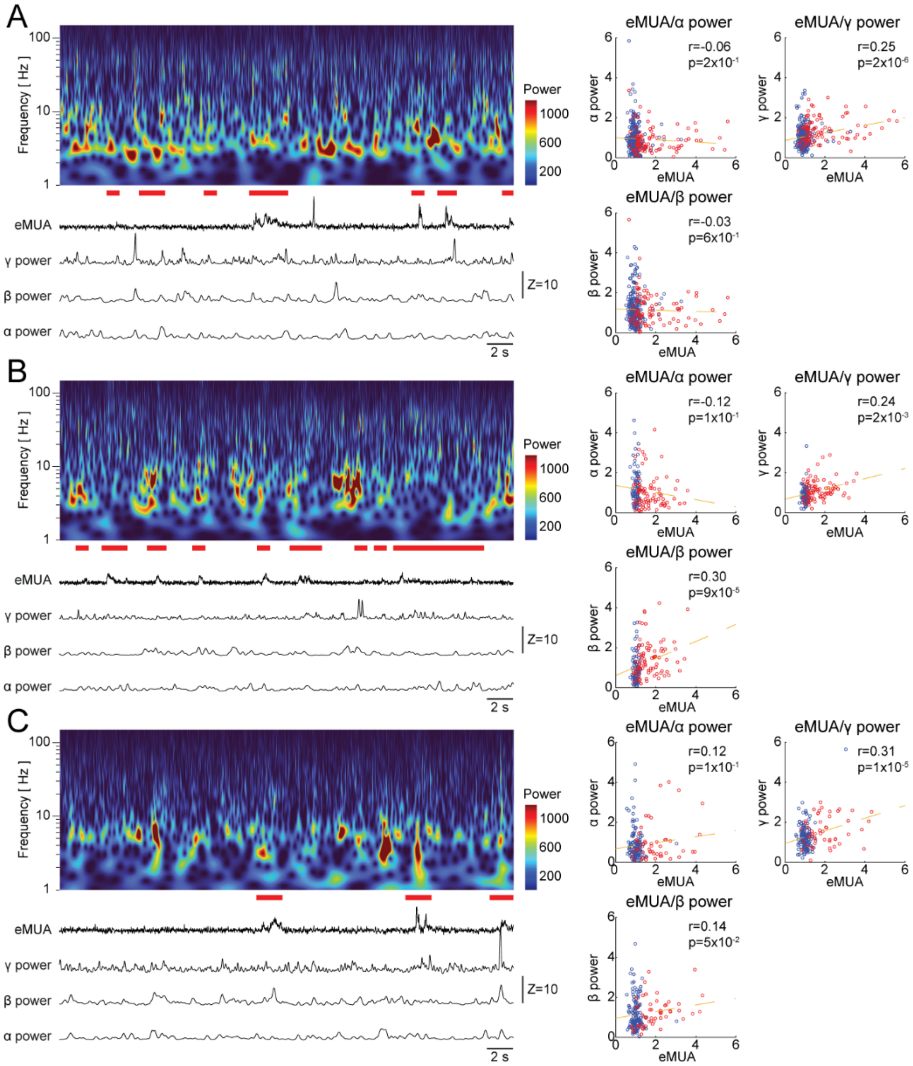

### Supplementary Figure 5

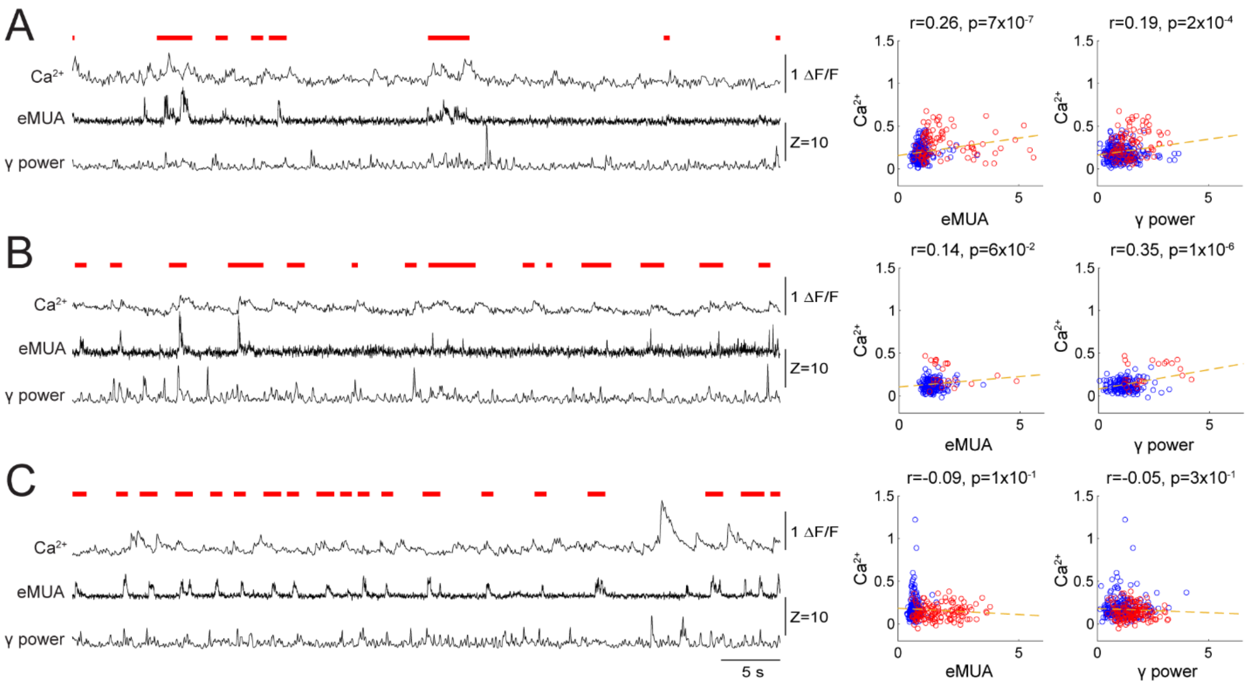

### Supplementary Figure 6

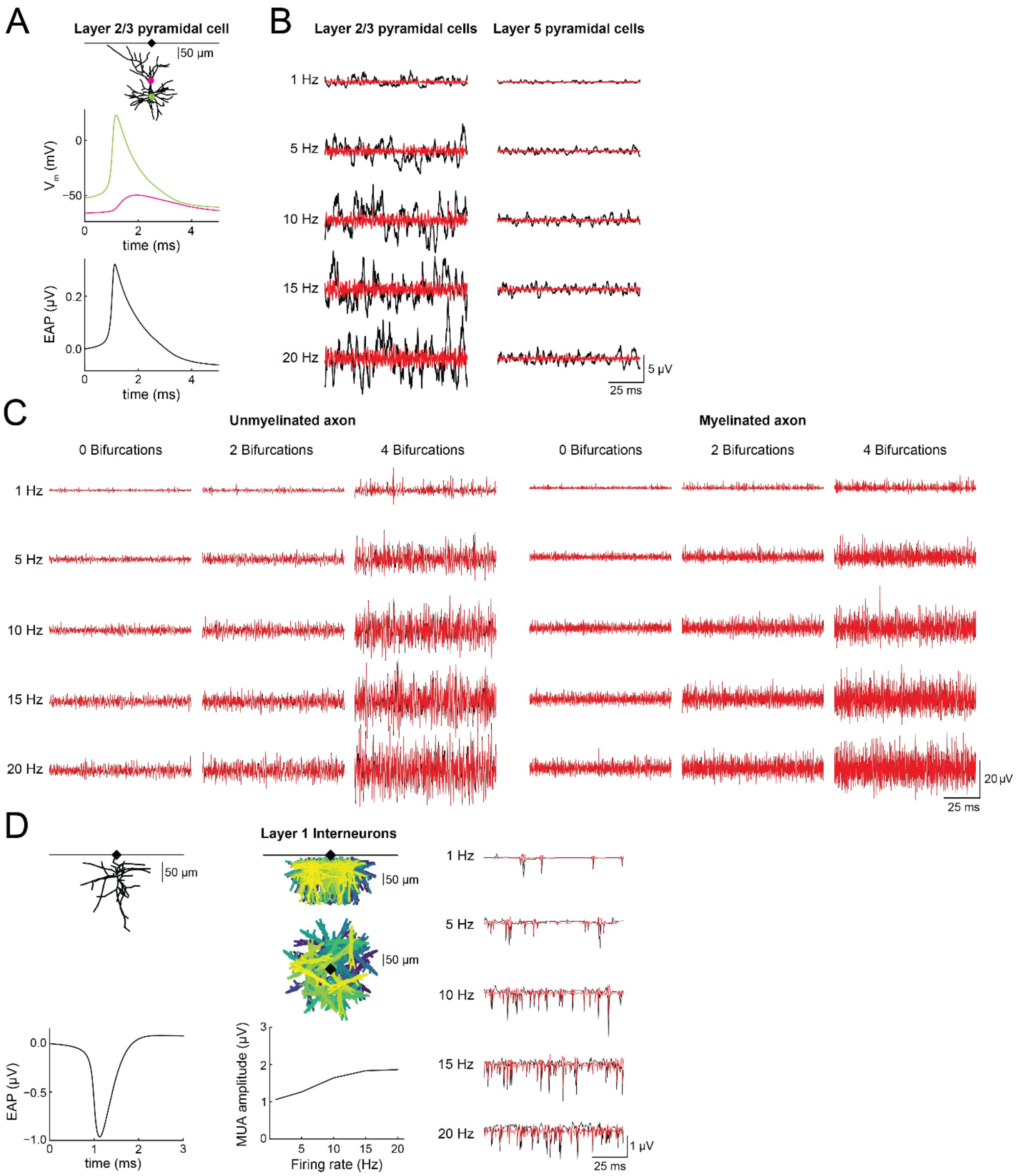
